# Lithium Restores Inhibitory Function and Neuronal Excitability through GSK-3β Inhibition in a Bipolar Disorder-Associated *Ank3* Variant Mouse Model

**DOI:** 10.1101/2023.10.26.564203

**Authors:** René N. Caballero-Florán, Kendall P. Dean, Andrew D. Nelson, Lia Min, Paul M. Jenkins

## Abstract

Bipolar disorder (BD) is a prevalent psychiatric condition characterized by mood dysregulation, psychosocial impairment, and an increased risk of suicide. The gene *ANK3* has been identified as a risk locus for BD through multiple genome-wide association studies (GWAS). However, the mechanisms by which *ANK3* variants influence BD pathophysiology and treatment response remain unclear. *ANK3* encodes ankyrin-G, a protein that organizes the axon initial segment (AIS) and nodes of Ranvier by scaffolding ion channels and cell adhesion molecules to the cytoskeleton. Recent studies show that ankyrin-G interacts with the GABA_A_ receptor-associated protein (GABARAP) to stabilize inhibitory synapses, potentially linking *ANK3* variants to inhibitory (GABAergic) signaling deficits associated with BD. We previously demonstrated that the BD-associated variant, *ANK3* p.W1989R, disrupts the ankyrin-G/ GABARAP interaction, resulting in inhibitory deficits and cortical pyramidal neuron hyperexcitability in mice. In this study, we investigate how lithium, a common BD therapeutic, modulates neuronal excitability in this model. Our findings show that chronic lithium treatment selectively enhances presynaptic GABAergic neurotransmission, reduces neuronal hyperexcitability, and partially rescues AIS length, without altering the density of GABAergic synapses. We also show that the selective glycogen synthase kinase-3 beta (GSK-3β) inhibitor Tideglusib recapitulates the enhancement of presynaptic GABAergic signaling. These findings shed new light on how *ANK3* variants may contribute to inhibitory deficits in BD and demonstrate that lithium treatment is able to restore these deficits, likely through GSK-3β inhibition. Furthermore, these findings highlight GSK-3β inhibition as a promising therapeutic strategy for treating BD and other neurological disorders affected by GABAergic dysfunction.

## INTRODUCTION

Bipolar disorder (BD) is a debilitating neuropsychiatric condition that affects approximately 2% of the global population (Merikangas et al., 2011). It is characterized by recurrent episodes of mania or hypomania and depression, marked by significant alterations in mood and behavior (American Psychiatric Association, 2013; Organization, 1993). In addition to mood disturbances, BD is associated with cognitive and functional impairments that substantially reduce quality of life for affected individuals (Cullen et al., 2016; Oldis et al., 2016). BD is also linked to premature mortality, largely due to comorbidities such as cardiovascular disease (Roshanaei-Moghaddam and Katon, 2009) and an increased risk of suicide (Gonda et al., 2012; Schaffer et al., 2015).

Despite the severity and prevalence of the disorder, the mechanisms underlying BD development are not fully understood. Evidence suggests the etiology of the disorder involves a combination of genetic, environmental, and neurochemical factors, leading to abnormalities in neuroanatomical structure and neuronal signaling (Carvalho et al., 2020). The heritability of BD is estimated to be as high as 85%, underscoring the significant role of genetics in susceptibility (Smoller and Finn, 2003). Several genome-wide association studies (GWAS) have identified the gene *ANK3* as a significant risk locus for the disorder (Ferreira et al., 2008; Lee et al., 2011; Muhleisen et al., 2014; Mullins et al., 2021; Psychiatric, 2011; Stahl et al., 2019). However, the precise mechanisms through which BD-associated *ANK3* variants contribute to the disorder and influence patient responses to BD therapeutics remain unclear.

*ANK3* encodes the protein ankyrin-G, which plays a critical role in the formation of excitable membrane domains in neurons (Bennett and Lorenzo, 2013; Nelson and Jenkins, 2017; Stevens and Rasband, 2021). Ankyrin-G mediates the assembly and function of the axon initial segment (AIS) and nodes of Ranvier by scaffolding voltage-gated Na+ and K+ channels to facilitate action potential (AP) generation and propagation in these regions (Jenkins and Bender, 2025; Nelson and Jenkins, 2017; Smith and Penzes, 2018; Stevens and Rasband, 2022). Additionally, ankyrin-G stabilizes inhibitory (GABAergic) synapses at the AIS and soma of neurons by interacting with the GABA_A_ receptor-associated protein (GABARAP), preventing endocytosis of GABA_A_ receptors (Nelson et al., 2020; Tseng et al., 2015).

Inhibitory (GABAergic) signaling is important for maintaining homeostasis of neuronal circuits and proper brain function, and deficits in GABAergic signaling have been implicated across multiple psychiatric disorders, including BD (Fatemi et al., 2013; Fatemi et al., 2017; Ozerdema et al., 2013; Schubert et al., 2015; Torrey et al., 2005). It is unknown whether common BD therapeutics, including lithium treatment, can reverse these deficits in inhibitory signaling. Lithium has been used in mood disorder treatment since its discovery in the mid-20th century (Shorter, 2009), and despite its limitations and side effects (Gitlin, 2016), it remains the gold standard for BD treatment (Fountoulakis et al., 2017; Yatham et al., 2018). Lithium monotherapy is effective for treating and preventing mania (Fountoulakis et al., 2022; Nestsiarovich et al., 2022) and is associated with anti-suicidal effects (Cipriani et al., 2013). Although lithium has been used as a therapeutic agent for over 60 years, the exact mechanisms underlying its effects remain under investigation.

Lithium has been shown to regulate a variety of cellular pathways involved in synaptic plasticity, brain circuitry, and neurotransmission (Bortolozzi et al., 2024). Many of these effects are thought to be mediated through lithium’s role as a glycogen synthase kinase-3 beta (GSK-3β) inhibitor (Bortolozzi et al., 2024). GSK-3β is a critical regulator of cellular homeostasis, and its dysregulation has been implicated in several disorders, including BD (Beurel et al., 2015; Jope and Johnson, 2004). Lithium inhibits GSK-3β both directly, by occupying a magnesium binding site on the enzyme, and indirectly, by increasing GSK-3β phosphorylation (Bortolozzi et al., 2024). While lithium treatment has been shown to regulate various cellular pathways through this mechanism, its effects on GABAergic signaling in the context of BD remain poorly understood.

Previously, we characterized a mouse model harboring the *Ank3* p.W1989R variant, originally identified in a family with BD, which abolishes the interaction between ankyrin-G and GABARAP (Nelson et al., 2020). *Ank3* p.W1989R mice display a significant reduction in the number of GABAergic synapses on the soma and AIS of cortical pyramidal neurons, as well as decreased forebrain gamma oscillations, indicative of reduced inhibitory function (Nelson et al., 2020; Tseng et al., 2015). Cortical pyramidal neurons from these mice also exhibit deficits in GABAergic neurotransmission, marked by a reduction in the frequency and amplitude of inhibitory post synaptic currents (IPSCs) (Nelson et al., 2020). This loss of inhibitory tone results in an increase in AP firing frequency and shortened AIS length, indicative of pyramidal neuron hyperexcitability (Nelson et al., 2020). Collectively, these results suggest that this *ANK3* variant may contribute to BD etiology through disrupted GABAergic signaling. Additionally, this raises the possibility that BD therapeutics, including lithium treatment, may function to restore these deficits and mitigate neuronal hyperexcitability.

This study investigates whether lithium treatment can restore GABAergic function and neuronal excitability in our previously described mouse model harboring the *Ank3* p.W1989R BD-associated variant. We demonstrate that chronic lithium treatment restores the frequency, but not the amplitude, of inhibitory postsynaptic currents (IPSCs) in *Ank3* p.W1989R mice. Lithium treatment also does not affect the density of GABAergic synapses, suggesting that it selectively enhances presynaptic GABAergic neurotransmission without affecting post-synaptic receptors in this mouse model. This enhancement of presynaptic GABA release is sufficient to normalize pyramidal neuron firing frequency and partially rescue AIS length abnormalities, indicating an overall reduction in neuronal hyperexcitability. Furthermore, the specific GSK-3β inhibitor Tideglusib mimicked the effects of chronic lithium treatment, enhancing presynaptic GABAergic neurotransmission without impacting postsynaptic properties, supporting a role of GSK-3β inhibition in modulating GABAergic activity. in contrast to chronic lithium treatment, the effects of Tideglusib were found to be AP-independent, suggesting differences between lithium therapy and specific GSK-3β inhibition.

This study demonstrates that chronic lithium treatment restores presynaptic GABAergic signaling and rescues neuronal hyperexcitability in *Ank3* p.W1989R mice likely through GSK-3β inhibition, highlighting GSK-3β inhibition as a promising therapeutic target for restoring neuronal function in BD and other mood disorders characterized by GABAergic dysfunction.

## RESULTS

### Chronic lithium treatment does not affect the density of GABAergic synapses on the soma or axon initial segment (AIS) of Ank3 p.W1989R cortical pyramidal neurons

GABAergic interneurons are essential for synchronizing neural networks and mediating higher-order cognitive functions in the brain (Buzsaki and Wang, 2012; Tamas et al., 2000). Disruptions in GABAergic signaling have been implicated in a variety of neuropsychiatric disorders, including BD (Fatemi et al., 2013; Fatemi et al., 2017; Ozerdema et al., 2013; Schubert et al., 2015; Torrey et al., 2005). Previously, we demonstrated that *Ank3* p.W1989R mice exhibit a significant reduction in the number of GABAergic synaptic clusters on the soma and AIS of cortical pyramidal neurons (Nelson et al., 2020). To investigate whether chronic lithium treatment can reverse these deficits, we performed immunohistochemistry on coronal brain sections from wild type (WT), *Ank3* p.W1989R (WR), and *Ank3* p.W1989R mice treated with lithium for 21 days (WR + Li). 21 days of lithium treatment was previously shown to rescue a subset of behavioral deficits in *Ank3* null mice (Zhu et al., 2017). *Ank3* p.W1989R littermates given lithium reached plasma lithium levels of: 1.07±0.039 mmol/L after treatment, concentrations similar to those recommended for patients undergoing lithium therapy (Nolen et al., 2019). Inhibitory synapses were identified by staining for the vesicular GABA transporter (vGAT), while ankyrin-G was used label the AIS (Fig. 1A). As expected, *Ank3* p.W1989R mice displayed a significant reduction in the density of GABAergic synaptic clusters on both the soma and AIS of pyramidal neurons (Fig. 1B, C) (Nelson et al., 2020). However, chronic lithium treatment did not restore the density of inhibitory synapses in these regions (Fig. 1). These findings prompted us to explore whether lithium treatment can affect other aspects of inhibitory neurotransmission in this mouse model, independent of changes in GABAergic synapse density.

**Figure 1.**
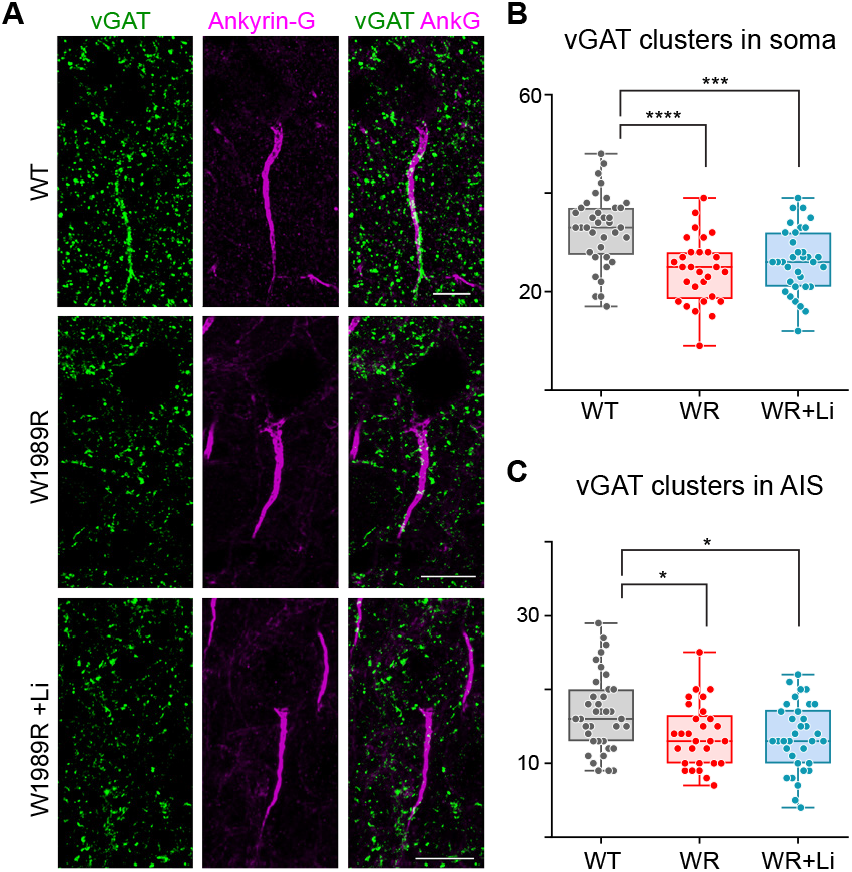
Lithium treatment does not affect the density of GABAergic synapses on the soma or axon initial segment (AIS) of cortical pyramidal neurons in *Ank3* p.W1989R mice. (A) Representative images of GABAergic synapses on pyramidal neurons in layer II/III somatosensory cortex of wild type (WT), *Ank3* p.W1989R (W1989R), and *Ank3* p.W1989R mice following 21 days of lithium treatment (W1989R+Li). Coronal brain sections were immunostained with a presynaptic GABAergic marker vGAT (green) and total ankyrin-G (magenta). Scale bars: 10 µm. (B) Quantification of the total number of vGAT-positive clusters per soma above a set intensity threshold from WT (gray), *Ank3* p.W1989R (WR) (red), and lithium treated *Ank3* p.W1989R (WR+Li) (blue). (C) Quantification of AIS vGAT-positive clusters from WT (gray), WR (red), and WR+Li (blue). One-way ANOVA: ****P < 0.0001, ***P = 0.0004 and P = ns (WT: n=37; WR: n=29; WR+Li: n=34).

### Chronic, but not acute, lithium treatment increases the frequency of spontaneous inhibitory postsynaptic currents (sIPSCs) in *Ank3* p.W1989R mice

To determine if lithium treatment rescues GABAergic function in *Ank3* p.W1989R mice, we performed whole-cell patch-clamp recordings on cortical sections from WT, *Ank3* p.W1989R, and *Ank3* p.W1989R mice treated with lithium for 19-21 days. Consistent with previous findings, both the frequency and amplitude of spontaneous inhibitory postsynaptic currents (sIPSCs) were reduced in cortical pyramidal neurons in *Ank3* p.W1989R mice compared to WT mice (Fig. 2A-C) (Nelson et al., 2020). After 21 days of lithium treatment, the amplitude of sIPSCs remained unchanged in *Ank3* p.W1989R mice (Fig. 2A-C). However, the frequency of sIPSCs in *Ank3* p.W1989R mice was rescued to levels observed in WT counterparts (Fig. 2A-C), reflecting an increase in presynaptic GABA release from cortical interneurons following chronic lithium treatment. WT mice treated with chronic lithium showed no significant changes in the frequency or amplitude of sIPSCs, indicating that lithium does not increase presynaptic GABA release in WT mice as it does in *Ank3* p.W1989R mice (Fig. S1). Furthermore, this effect is dependent on chronic lithium administration, as acute lithium exposure did not rescue sIPSC frequency in *Ank3* p.W1989R mice (Fig. S2). These results indicate that chronic, but not acute, lithium treatment selectively enhances presynaptic GABAergic transmission in this mouse model.

**Figure 2.**
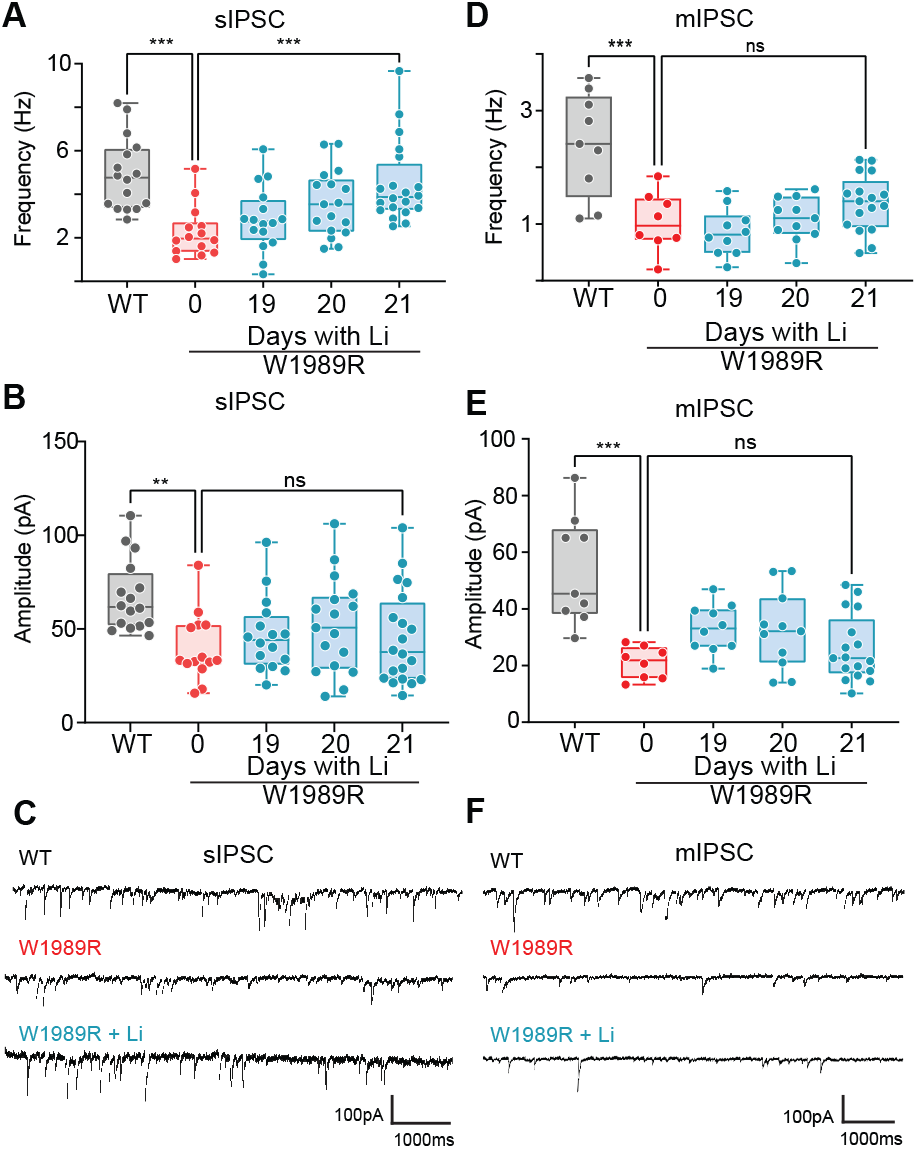
Chronic lithium treatment increases IPSC frequency, while not affecting current amplitude, in *Ank3* p.W1989R mice. (A) Quantification of sIPSC and frequency (Hz) and (B) amplitude (pA) from cortical neurons from brain slices of wild type (WT) (gray), *Ank3* p.W1989R (W1989R) (red), and *Ank3* p.W1989R (W1989R) after chronic lithium treatment 19-21 days (blue). Kruskal-Wallis, Dunn‘s multiple comparisons test ***P < 0.01 and P= ns (WT: n=16; W1989R: n = 15; W1989R/Day 19 + Li: n = 16; W1989R/Day 20 + Li: n = 17 and W1989R/Day 21 + Li n = 20). (C) Representative traces of sIPSC plots (A) and (B). (D) Quantification of mIPSC frequency (Hz) and amplitude (E) of WT (gray), W1989R (red), and W1989R after chronic lithium treatment 19-21 days (blue). Ordinary one-way ANOVA, Tukey‘s multiple comparisons tests: ***P < 0.001, and P = ns. (WT: n = 9; W1989R: n = 8; W1989R/Day 19 + Li: n = 10; W1989R/Day 20 + Li: n = 11 and W1989R/Day 21 + Li n = 17). (F) Representative traces of mIPSC in (D) and (E).

### Chronic lithium treatment does not alter miniature inhibitory postsynaptic currents (mIPSCs) in *Ank3* p.W1989R mice

To examine whether alterations in GABAergic transmission following chronic lithium treatment are AP-independent, we measured mIPSCs in the presence of TTX. Excitatory pyramidal neurons in *Ank3* p.W1989R cortical sections exhibited significantly reduced mIPSC frequency and amplitude compared to those from WT, consistent with previous findings (Fig. 2D-F) (Nelson et al., 2020). Interestingly, chronic lithium treatment did not restore the frequency or amplitude of mIPSCs in *Ank3* p.W1989R mice (Fig. 2D-F), indicating that lithium’s effects on GABAergic transmission are dependent on AP firing in this model.

WT mice treated with chronic lithium also showed no significant changes in the frequency or amplitude of mIPSCs (Fig. S1). Additionally, no sex-related differences were observed following lithium treatment (Fig. S3). Collectively, these results suggest that chronic, but not acute, lithium therapy selectively enhances presynaptic GABAergic transmission from interneurons in *Ank3* p.W1989R mice in an AP-dependent manner, without impacting the amplitude of postsynaptic currents recorded from pyramidal neurons.

### Tideglusib increases the frequency of sIPSCs and mIPSCs in *Ank3* p.W1989R mice

Inhibition of GSK-3β is a key mechanism thought to mediate lithium‘s therapeutic effects (Bortolozzi et al., 2024), and Tideglusib is an approved GSK-3β inhibitor (Dominguez et al., 2012; Fuchs et al., 2018; Morales-Garcia et al., 2012; Sereno et al., 2009). To determine if the effects of lithium treatment on presynaptic GABAergic function are recapitulated by specific GSK-3β inhibition, we treated *Ank3* p.W1989R mice with Tideglusib for 20 days and measured sIPSCs and mIPSCs in layer II/III cortical pyramidal neurons. Tideglusib treatment resulted in an increase in both sIPSC and mIPSC frequency in *Ank3* p.W1989R mice (Fig. 3A, C, D, F), with no impact on the amplitude of either sIPSCs or mIPSCs (Fig. 3B, C, E, F). These findings demonstrate that, like chronic lithium treatment, specific GSK-3β inhibition enhances presynaptic GABAergic neurotransmission in *Ank3* p.W1989R mice without altering postsynaptic receptor properties. Interestingly, unlike chronic lithium treatment, the effects of Tideglusib on inhibitory signaling appear to be AP-independent, as demonstrated by the increase in mIPSC frequency following Tideglusib treatment (Fig. 3). These results suggest that GSK-3β inhibition is likely an important mechanism underlying lithium’s modulation of presynaptic GABAergic neurotransmission. However, key differences exist between lithium and Tideglusib treatment, particularly regarding the AP-dependence of this process.

**Figure 3.**
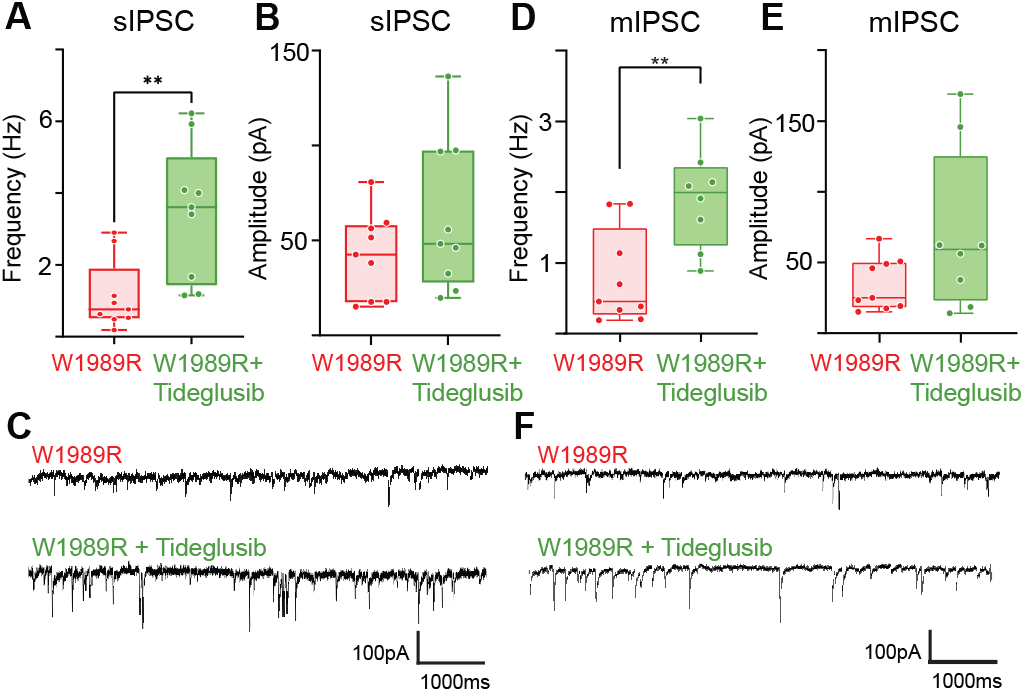
Chronic Tideglusib treatment increases sIPSC and mIPSC frequency while not affecting current amplitude in *Ank3* p.W1989R mice. (A) Quantification of sIPSC frequency (Hz) and (B) amplitude (pA) from cortical neurons from brain slices of Ank3 p.W1989R (W1989R) (red), and Ank3 p.W1989R (W1989R) with Tideglusib treatment (green). Unpaired t-test **P < 0.005 (W1989R + vehicle: n = 9; and W1989R + Tideglusib n = 9). (C) Representative traces of sIPSC in (A) and (B). (D) Quantification of mIPSC frequency (Hz) and (E) amplitude (pA) from cortical neurons from brain slices of Ank3 pW1989R (W1989R) (red), and Ank3 pW1989R (W1989R) chronic Tideglusib treatment (green). Unpaired t-test **P < 0.005 (W1989R + vehicle: n = 9; and W1989R + Tideglusib n = 9) (F) Representative traces of mIPSC plots (D) and (E).

### Chronic lithium treatment restores action potential firing rate and frequency in *Ank3* p.W1989R mice

To assess the impact of chronic lithium treatment on pyramidal neuron excitability, we compared evoked AP firing frequency in WT, *Ank3* p.W1989R, and *Ank3* p.W1989R mice treated with lithium for 21 days. Consistent with previous findings, *Ank3* p.W1989R mice exhibited significantly increased AP firing frequencies in response to escalating current injections, compared to WT mice (Fig. 4A, B) (Nelson et al., 2020). Additionally, the maximum firing frequency of *Ank3* p.W1989R neurons was significantly higher than that of WT neurons (Fig. 4C). Following chronic lithium treatment, both the firing rates across different current injections and the maximum firing frequency of APs were restored to levels observed in WT mice (Fig. 4A, B, C). Cell membrane and AP properties were not affected after lithium treatment (Fig.S4). While the threshold of the first AP generated from the AP train was significantly more depolarized in lithium-treated *Ank3* p.W1989R mice compared with WT mice, this difference was not seen in single evoked APs (Fig.S4). These data indicate that chronic lithium treatment restores pyramidal neuron excitability in *Ank3* p.W1989R to levels seen in WT mice without affecting single AP properties.

**Figure 4.**
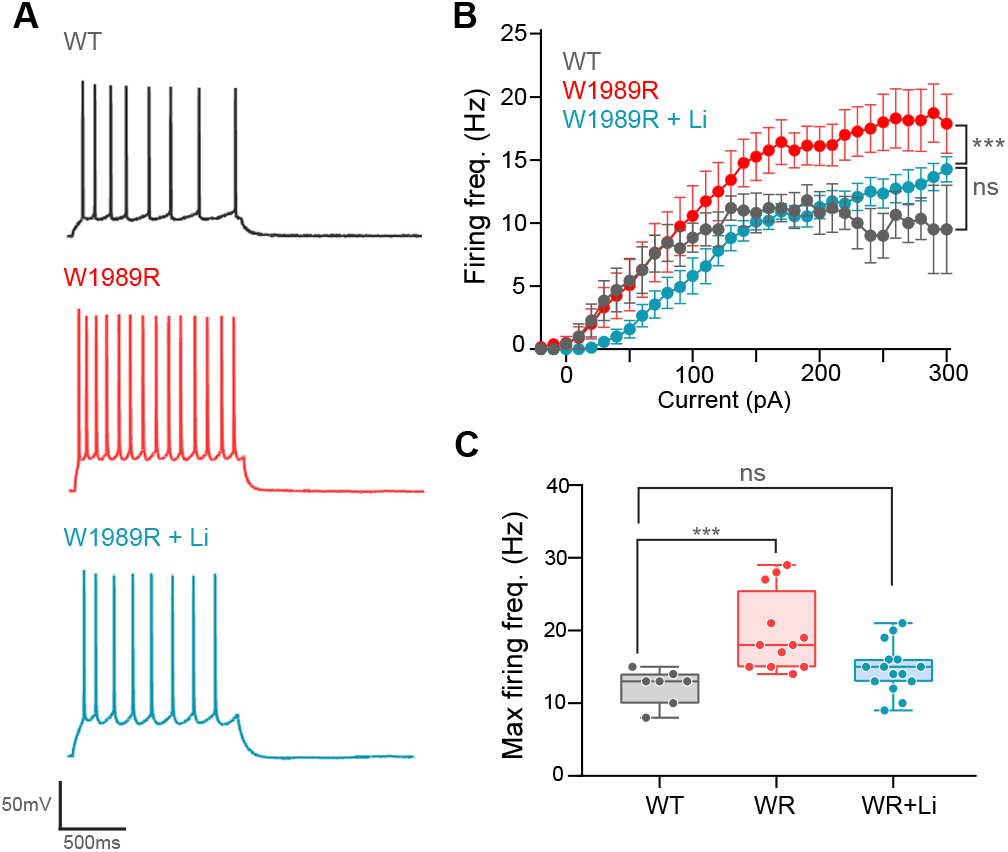
Pyramidal neuron firing frequency is restored in *Ank3* p.W1989R mice following chronic lithium treatment. (A) Representative traces of APs generated by current injection in wild type (WT) (gray), *Ank3* p.W1989R (WR) (red), and *Ank3* p.W1989R after lithium treatment (WR + Li) (blue) neurons. (B) Average firing rate (Hz) vs. current injection (pA) from evoked APs in cortical neurons from wild type (WT) (gray), *Ank3* p.W1989R (W1989R) (red), and *Ank3* p.W1989R after lithium treatment (W1989R + Li) (blue). Two-way ANOVA, Tukey‘s multiple comparisons tests: WT vs. W1989R ***P < 0.001, WT vs. W1989R + Lithium P = ns, and W1989R vs. W1989R + Lithium ***P < 0.001. (C) Maximum firing frequency (Hz) graph of pyramidal neurons from WT (gray), *Ank3* p.W1989R (WR) (red), and *Ank3* p.W1989R after lithium treatment (WR + Li) (blue). Kruskal-Wallis, Dunn‘s multiple comparisons tests: WT vs. W1989R ***P < 0.001, WT vs. W1989R + Lithium P = ns, W1989R + Lithium vs W1989R P < 0.05 (WT: n = 7; W1989R: n = 12; W1989R + Li: n = 15).

### Chronic lithium treatment partially restores AIS length deficits in *Ank3* p.W1989R mice

The axon initial segment (AIS) is dynamically regulated in response to neuronal activity, and alterations in AIS morphology can reflect changes in neuronal signaling and network activity (Huang and Rasband, 2018). Excitatory pyramidal neurons from *Ank3* p.W1989R mice exhibit a shortened AIS, which is thought to serve as a compensatory mechanism for neuronal hyperexcitability (Nelson et al., 2020). To assess whether lithium treatment impacts AIS morphology, we quantified the length of the AIS in layer II/III cortical pyramidal neurons from WT, *Ank3* p.W1989R, and *Ank3* p.W1989R mice treated with lithium for 21 days. Ankyrin-G immunostaining was used to identify the AIS in each sample. Interestingly, lithium treatment resulted in a significant increase in AIS length in *Ank3* p.W1989R mice, although it remained shorter than that of WT mice (Fig. 5A-B). This partial restoration of AIS length may reflect lithium‘s ability to mitigate neuronal hyperexcitability in *Ank3* p.W1989R mice, thereby reducing the need for compensatory mechanisms such as AIS shortening.

**Figure 5.**
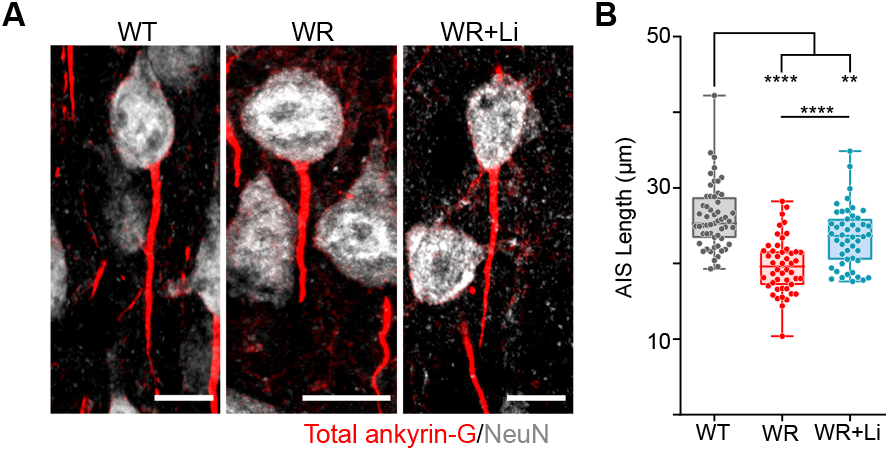
Lithium treatment increases the AIS length in *Ank3* p.W1989R mice. (A) Representative images of pyramidal neuron AIS length from layer II/III somatosensory cortex of wild type (WT), *Ank3* p.W1989R (WR), and *Ank3* p.W1989R following lithium treatment (WR+Li). Immunostaining for total ankyrin-G (red) and NeuN (white). Scale bars: 10 µm. (B) Quantification of AIS length between WT (gray), WR (red), and WR+Li (blue). One-way ANOVA: ****P < 0.0001, ****P < 0.0001 and **P = 0. 0033 (WT: n=55; WR: n=48; WR+Li: n=48).

## DISCUSSION

Bipolar disorder (BD) is a prevalent psychiatric disorder, characterized by behavioral dysregulation and elevated risk of suicide. The gene *ANK3*, encoding ankyrin-G, has been identified as a significant risk locus for BD through multiple genome-wide association studies (GWAS). However, the precise mechanisms by which variants in *ANK3* contribute to BD pathophysiology, as well as their influence on treatment response, remain incompletely understood. Previously, we demonstrated that the *ANK3* p.W1989R variant, identified in a BD patient cohort, leads to impaired inhibitory signaling and neuronal hyperexcitability in mice. In the present study, we investigate how chronic lithium treatment modulates neuronal excitability and inhibitory signaling in this model, finding that it selectively enhances presynaptic inhibitory signaling and alleviates several hallmarks of neuronal hyperexcitability. Specifically, we show that chronic lithium treatment in *Ank3* p.W1989R mice selectively augments presynaptic interneuron-mediated GABAergic neurotransmission, reduces pyramidal neuron hyperexcitability, and partially restores abnormalities in AIS length, without affecting GABAergic synapse density. We also demonstrate that treatment with the selective glycogen synthase kinase-3 beta (GSK-3β) inhibitor, Tideglusib, produces similar results, enhancing presynaptic GABAergic neurotransmission. These findings suggest that both chronic lithium treatment and selective GSK-3β inhibition may exert therapeutic effects in BD patients by enhancing interneuron-mediated GABAergic signaling, thus mitigating neuronal hyperexcitability.

Our results demonstrate that chronic lithium treatment in *Ank3* p.W1989R mice selectively increases the frequency, but not amplitude, of sIPSCs. This suggests that lithium treatment induces an increase in GABA release from presynaptic terminals without altering postsynaptic receptor properties. Interestingly, the GSK-3β inhibitor Tideglusib produced similar effects, enhancing the frequency of sIPSCs. This is likely due to the shared mechanism of GSK-3β inhibition between lithium and Tideglusib. GSK-3β is a key regulator of synaptic plasticity and neuronal signaling, and its inhibition has been shown to exert therapeutic effects in the treatment of BD and other mood disorders (Bortolozzi et al., 2024). These findings suggest that the effects of lithium on GABAergic signaling in *Ank3* p.W1989R mice are likely mediated, at least in part, by GSK-3β inhibition. Furthermore, they reinforce the growing body of evidence that supports GSK-3β as a critical target in BD treatment and underscore its potential as a therapeutic pathway for modulating GABAergic function in BD.

While both chronic lithium and Tideglusib treatments restored sIPSC frequency, only Tideglusib was able to rescue deficits in mIPSC frequency observed in *Ank3* p.W1989R mice. This is particularly noteworthy, as, unlike sIPSCs, mIPSCs are measured in the presence of the sodium channel inhibitor TTX, which blocks AP firing. While both lithium and Tideglusib inhibit GSK-3β, the differences in their effects on AP-dependent versus AP-independent processes may be attributed to the broader spectrum of effects lithium has on neuronal excitability, whereas Tideglusib’s more selective inhibition of GSK-3β likely produces more localized changes in neurotransmission that are independent of AP firing. Neither chronic lithium nor Tideglusib treatment was able to restore the amplitude of mIPSCs, reinforcing the notion that neither is capable of modulating postsynaptic receptor properties in this mouse model.

The density and function of postsynaptic inhibitory receptors influence the amplitude of inhibitory post-synaptic currents (IPSCs). The *Ank3* p.W1989R mutation disrupts the interaction between ankyrin-G and GABARAP, which results in enhanced endocytosis of the GABA_A_ receptor and a reduction in the number of inhibitory synapses (Nelson et al., 2020; Tseng et al., 2015). Gephyrin, one of the main scaffolding proteins involved in the stabilization of inhibitory synapses, is phosphorylated by GSK-3β, and inhibition of GSK-3β through lithium treatment has been shown to increase gephyrin clustering at inhibitory synapses (Tyagarajan et al., 2011). While the exact relationship between gephyrin and ankyrin-G at inhibitory synapses remains unclear, an increase in gephyrin clustering through GSK-3β inhibition could potentially stabilize GABA synapses to overcome deficits associated with *Ank3* p.W1989R. However, we find that lithium treatment is unable to restore deficits in GABAergic synapse density in this mouse model. This observation raises further questions about the relationship between these scaffolding proteins and their role in stabilizing GABAergic synapses, as these findings suggest that increased gephyrin clustering alone cannot restore the number of inhibitory synapses, at least in this *Ank3* p.W1989R variant model.

Several studies have explored the impact of lithium treatment on *Ank3*-deficient mice (Garza et al., 2018; Gottschalk et al., 2017; Piguel et al., 2023). While this study provides compelling evidence for the role of chronic lithium treatment in modulating GABAergic neurotransmission and neuronal excitability in *Ank3* p.W1989R mice, it is important to note that mice do not fully recapitulate all aspects of BD pathology. Future studies employing human-derived model systems could help bridge the gap between animal models and BD patients. Additionally, sleep disruption is a common phenotype in many neuropsychiatric disorders, including BD, and abnormal sleep patterns have been observed in mice harboring a BD-associated *Ank3-1b* deletion variant (Villacres et al., 2023). Characterizing sleep abnormalities and their potential mediation by lithium treatment in this mouse model could offer further insights into how lithium therapies influence BD patient behavior. Furthermore, while GSK-3β inhibition appears to play a key role in modulating GABAergic transmission, other lithium-sensitive signaling pathways may also contribute to its therapeutic effects and should be explored in future studies.

Our results demonstrate that chronic lithium treatment partially restores inhibitory deficits, normalizing the excitability of cortical pyramidal neurons, without altering the number of inhibitory synapses in a BD-patient *ANK3* variant mouse model. Furthermore, we find that this restoration of excitatory and inhibitory balance is, at least in part, due to the inhibition of GSK-3β, a key mediator of lithium’s therapeutic action. These findings provide new insights into how lithium treatment may enhance inhibitory signaling and promote therapeutic effects in BD patients, contributing to the growing body of research implicating GSK-3β inhibition as a primary mechanism underlying lithium’s mood-stabilizing effects. By further investigating how GSK-3β inhibition enhances inhibitory signaling and integrating mouse studies with BD patient-derived neurons, we may be able to develop more targeted and effective therapeutic strategies for BD that not only address the symptoms but also specifically restore the underlying neuronal deficits.

## MATERIALS AND METHODS

### Animals

*Ank3* p.W1989R mice were described previously (Nelson et al., 2020). *Ank3* p.W1989R heterozygous male and female mice were crossed to generate respective genotypes and littermates were used for all experiments. Animals were group housed (2 to 5 per ventilated cage) with 24-hour (12/12 dark/light cycle) access to food and water. Litters from the heterozygous breeders were weaned at postnatal day 21 (P21) and raised to P42-48. For mice that underwent lithium therapy, lithium chow was provided at P21, while control mice were fed with regular chow (vehicle), as described below. All experimental procedures were performed with the guidelines for animal care of the institutional animal care and use committee (IACUC), and university laboratory animal management (ULAM) at the University of Michigan.

### Pharmacological treatments and measurement of plasma lithium levels

Littermates of both sexes were fed with regular chow or 0.2% (g/kg chow) lithium carbonate diet (Harlan Teklad TD.170313/12) for 14 days followed by 7 days of 0.36% (g/kg chow) to achieve lithium therapeutic levels as described previously (Zhu et al., 2017). During lithium treatment, each cage of mice was supplemented with an extra bottle of 0.9% NaCl to reduce the risk of dehydration and the bedding was changed daily. For Tideglusib treatment, mice were administrated Tideglusib (Sigma-Aldrich, St. Louis, MO, USA) or vehicle (corn oil; Sigma-Aldrich, St. Louis, MO, USA) via subcutaneous injection every other day for 20 days (20 mg/kg body weight) (Fuchs et al., 2018). Electrophysiology recordings were carried out the next day following cessation of treatment. Blood was collected immediately after euthanasia and saved in tissue plasminogen activator (TPA)/heparin (100 µL). Samples were sent to the University of Michigan clinical core to measure serum lithium concentration.

### *In vitro* electrophysiology recordings

Mice were anesthetized with isoflurane and decapitated (all procedures are by approved UM IACUC protocol PRO00010191). The brain was quickly removed from the skull and placed in 4°C slicing solution (in mM): 62.5 NaCl, 2.5 KCl, 1.25 KH_2_PO_4_, 26 NaHCO_3_, 5 MgCl_2_, 0.5 CaCl_2_, 20 glucose and 100 sucrose (pH = 7.4 with O_2_/CO_2_ 95/5%, ∼315 mOsm). The location of the somatosensory cortex in coronal brain slices was identified according to the mouse brain atlas by Paxinos and Franklin(Paxinos and Franklin, 2019), and coronal slices (∼300µm thick) containing somatosensory cortical layers II/III were cut on a microtome (VF-300, CompresstomeTM; Precisionary instruments, Natick MA). All recordings were taken from the same brain region.

Brain slices were transferred to a chamber maintained at room temperature in artificial cerebrospinal fluid (ACSF) (in mM): 125 NaCl, 2.5 KCl, 1.25 KH_2_PO_4_, 26 NaHCO_3_, 1 MgCl_2_, 2 CaCl_2,_ and 20 glucose, pH 7.4 and ∼300 mOsm, for at least 1 hour before recording. Individual slices were transferred to a recording chamber perfused with ACSF (1–2 mL/min). Recording micropipettes were pulled for a final resistance of 5-7 MΩ (P-97; Sutter Instruments, Novato, CA) from borosilicate glass capillaries (1.5 mm O.D.; Harvard Apparatus, Holliston, MA) and filled with solution (in mM): 135 K-Gluconate, 4 NaCl, 0.4 GTP, 2 Mg-ATP,0.5 CaCl_2_, 5 EGTA and 10 HEPES, pH 7.25, ∼290 mOsm. The neurons were identified optically using a Nikon Eclipse FN-1 microscope with a 40X water-immersion objective and a DAGE-MTI IR-1000 video camera. Neurons were characterized using IR-DIC to evaluate their orientation, morphology, and spiking properties. Signals were recorded with Axoclamp 700B amplifier (Axon Instruments, Union City, CA), low pass filtered at 10 kHz. Whole-cell patch-clamp recordings were performed on cells with a high cell resistance (>8 GΩ before break-in). Current clamp recordings were obtained and analyzed from neurons in layers II/III of somatosensory cortex as previously reported (Nelson et al., 2020). The neurons were recorded by applying negative and positive current pulses of 10 pA for 1000 ms to calculate the rheobase (minimum current to fire the first AP) and the maximum firing frequency response, and single positive pulses of 1 ms to measure single AP properties as previously described (Nelson et al., 2020). For GABAergic currents, the K-gluconate in the internal solution was replaced with CsCl [140 mM], and the recordings were acquired at 2 kHz holding at -70 mV. Spontaneous inhibitory postsynaptic currents (sIPSCs) and miniature spontaneous inhibitory postsynaptic currents (mIPSCs) were pharmacologically isolated using the AMPA/kainite receptor antagonist 6-cyano-7-nitroquinoxaline-2,3-dione (CNQX) [10 µM], and the NMDA antagonist DL-2-Amino-5-phosphonopentanoic acid (DL-AP-5) [100 µM]. The use of CNQX and DL-AP-5 pharmacologically isolates sIPSCs by specifically blocking excitatory signals, thereby mitigating potential interference with the measurement of inhibitory currents. To assess mIPSCs, the perfusion was supplemented with 1 µM Tetrodotoxin (TTX) to inhibit synaptic responses reliant on action potentials. Continuous monitoring of access resistance was conducted during the experiment, and trials were terminated if changes exceeding 20% were observed. The analysis entailed assessing the frequency and amplitude of these events, as described earlier. The events were automatically detected using Minianalysis (Synaptosoft Inc.) and were visually examined to eliminate any potential erroneous noise (Nelson et al., 2020).

### Immunocytochemistry of brain sections

After 21 days of lithium treatment, P42 mice were administered a ketamine/xylazine mixture (80 mg/kg body weight ketamine and 10 mg/kg xylazine) via intraperitoneal injection. The mice were euthanized by cardiac perfusion of PBS followed by 4% paraformaldehyde and the brain was immediately removed and fixed overnight in 4% paraformaldehyde. The next day, the brains were processed using a standard single-day paraffin preparation protocol (PBS wash followed by dehydration through 70, 95, and 100% ethanol with final incubations in xylene and hot paraffin under vacuum) using a Leica ASP 300 paraffin tissue processor. Paraffin sections were cut at 7 μm thickness using a Leica RM2155 microtome and placed on glass slides. Sections were deparaffinized and rehydrated using a standard protocol of washes: 3 × 4-min xylene washes, 2 × 2-min 100% ethanol washes, and 1 × 2-min 95%, 80%, and 70% ethanol washes followed by at least 5 min in ddH_2_O. Antigen retrieval was then conducted by boiling the deparaffinized brain sections for 20 min in 10 µM sodium citrate in a microwave oven. Sections were cooled, washed for 15 min in ddH_2_O, rinsed in PBS for 5 min, and blocked using blocking buffer (5% BSA, 0.2% Tween 20 in PBS) for 1 hour at room temperature. Slides were incubated overnight at 4°C with primary antibodies diluted in a blocking buffer. On the following day, slides were washed 3 times for 15 min with PBS containing 0.2% Tween 20, incubated with secondary antibodies diluted in blocking buffer for 1 hour at room temperature, washed 3 times for 15 min, and mounted with Prolong Gold. Samples were imaged on a Zeiss LSM 880 with a 63X NA1.4 Oil/DIC Plan-Apochromat objective at 488, 561, and 647 nm lasers. AIS length and GABAergic synapse number were measured using maximum intensity projections of z-stacks. We only quantified AISs that were entirely contained within the bounds of the Z-stack. The number of vGAT-positive puncta was quantified above a set fluorescence intensity consistent across all samples. N refers to the number of mice used in the experiment, whereas n refers to the total number of neurons measured.

### Statistical Analysis

Statistical analysis was conducted using GraphPad Prism (Version 8.0) software. To assess the significance of differences between groups, Unpaired t-tests and ANOVAs were employed, with verification that the results adhered to the assumptions of normality and homogeneity of variances using the Shapiro-Wilk and Bartlett tests, respectively. We presented our statistical results by elucidating the specifics of the applied statistical tests, including the statistical value, degrees of freedom, and the exact p-value, tailored to the specific test used. In cases where the Gaussian distribution criterion was not met, the Kruskal-Wallis test with Dunn‘s post hoc analysis was applied. When inequality of variances was identified, Welch‘s ANOVA was used, followed by the post hoc Dunnett‘s test. In all instances, the criterion for statistical significance was set at Alpha (α) = 0.05. For tests with observations of statistical significance post-hoc power analyses were conducted using G Power (3.1). These analyses considered the alpha level, sample size based on the group average, and effect size to compute the achieved equivalent statistical power (1-beta (β) error probability) of 0.8 or higher.

## ACKNOWLEDGEMENTS

This work was supported by the One Mind Rising Star Bipolar Disorder Translational Research Award (PMJ), Cellular and Molecular Biology at Michigan T32GM007315 (KPD), NIH R21NS113022 (PMJ) and NIH R01MH126960 (PMJ). We thank the University of Michigan Clinical Core for measuring plasma lithium concentrations in mice after treatment. We thank Dr. Kathleen Ignatoski and Dr. Julie Gupta, for their assistance with lithium therapy and mouse husbandry in the early experimental stages. We thank the members of Dr. Lori Isom’s lab, including Drs. Luis Lopez-Santiago and Yukun Yuan from University of Michigan, for their help in the analysis, discussion, and interpretation of the obtained results. We also thank members of the Jenkins laboratory, Dr. Kevin Bender (UCSF), and Dr. Erica Levitt (University of Michigan) for their valuable feedback on this manuscript.

## AUTHOR CONTRIBUTIONS

RNFC and PMJ designed research. RNCF and ADN performed experiments and analyzed data. LM performed the genotyping of the mice. LM and KPD participated in the editing and final design of the manuscript.

## SUPPLEMENTARY MATERIALS

Figures S1 to S4

