## Supplemental Information for "Lithium Restores Inhibitory Function and Neuronal Excitability through GSK-3β Inhibition in a Bipolar Disorder-Associated *Ank3* Variant Mouse Model"

**Supplementary Material**

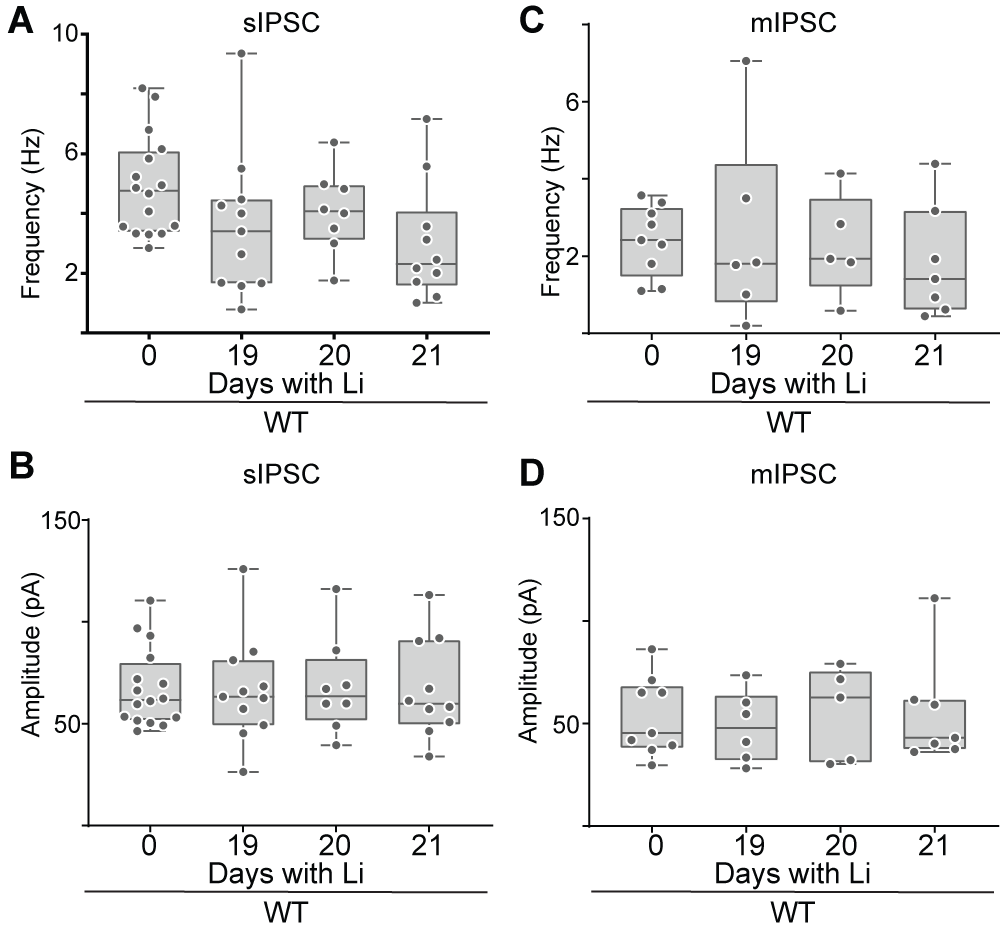

**Figure S1.** **Chronic lithium treatment does not affect IPSC frequency or amplitude in WT mice.** (A) Quantification of sIPSC frequency (Hz) from cortical neurons from brain slices of WT mice without treatment and after chronic lithium treatment 19-21 days: Ordinary one-way ANOVA (F (3, 41) = 2.326, P = 0.0889). (WT: n=16; WT/Day 19 + Li: n=12; WT/Day 20 + Li: n=8 and WT/Day 21 + Li n=10) (B) and amplitude (pA): Ordinary one-way ANOVA (F (3, 41) = 0.1093, P = 0.9542). (WT: n=16; WT/Day 19 + Li: n=11; WT/Day 20 + Li: n=8 and WT/Day 21 + Li n=10). (C) Quantification of mIPSC frequency (Hz) of WT (gray) and WT after chronic lithium treatment 19-21 days. Ordinary one-way ANOVA (F (3, 23) = 0.2659, P=0.8492) (D) and amplitude (pA): Ordinary one-way ANOVA (F (3, 41) = 0.5547, P=0.6479), (WT: n=9; WT/Day 19 + Li: n=6; WT/Day 20 + Li: n=5 and WT/Day 21 + Li n=7). N=3-6 mice.

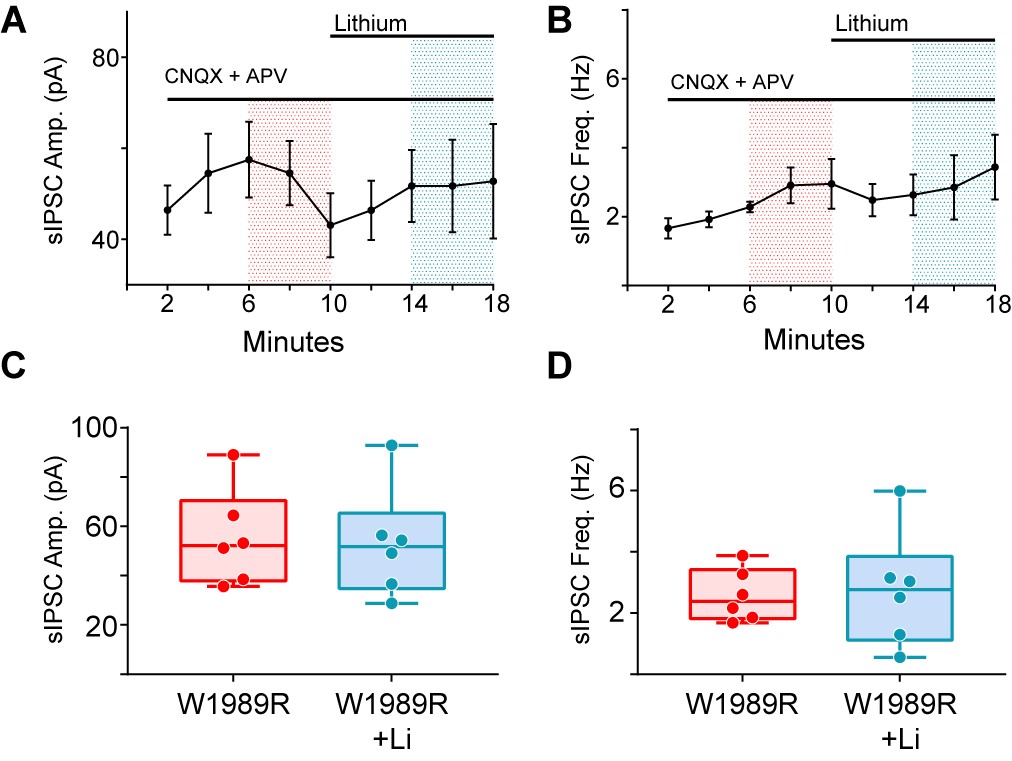

**Figure S2. Acute lithium treatment does not modify sIPSCs in layer II/III cortical neurons from *Ank3* p.W1989R mice.** (A) The temporal curve of sIPSC amplitude (pA) and (B) frequency (Hz) in response to acute lithium (0.8mmol/L) for 8 minutes. (C) Quantification of the sIPSC amplitude (pA) (T-test (t=0.1916, df=10, P= 0.852) in (A). Quantification of the sIPSC frequency (Hz) (T-test (t=0.2119, df=10, P= 0.836)) in (B). *Ank3* p.W1989R (W1989R) (red), and *Ank3* p.W1989R in the presence of lithium (W1989R +Li) (blue) in acute brain slices. (W1989R: n=6; W1989R + Li: n=6). For quantification in plots (A) and (C) the average from minutes 4-8 was estimated as a control and 14-18 minutes for the lithium condition. N=2 mice.

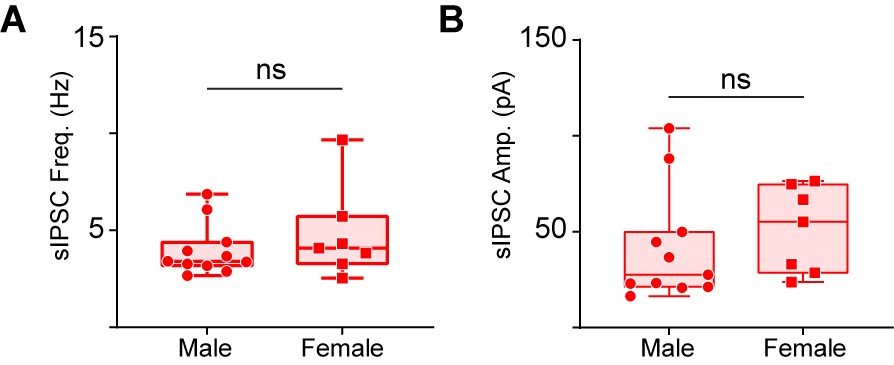

**Figure S3. Lithium treatment causes identical effects in male and female *Ank3* p.W1989R mice.** (A) sIPSC frequency (Hz) and (B) amplitude (pA) in response to acute lithium treatment for 21 days, in acute brain slices from Males (red circles) and females (red squares). Mann Whitney test P = ns (Males n=11 and Females n = 7).

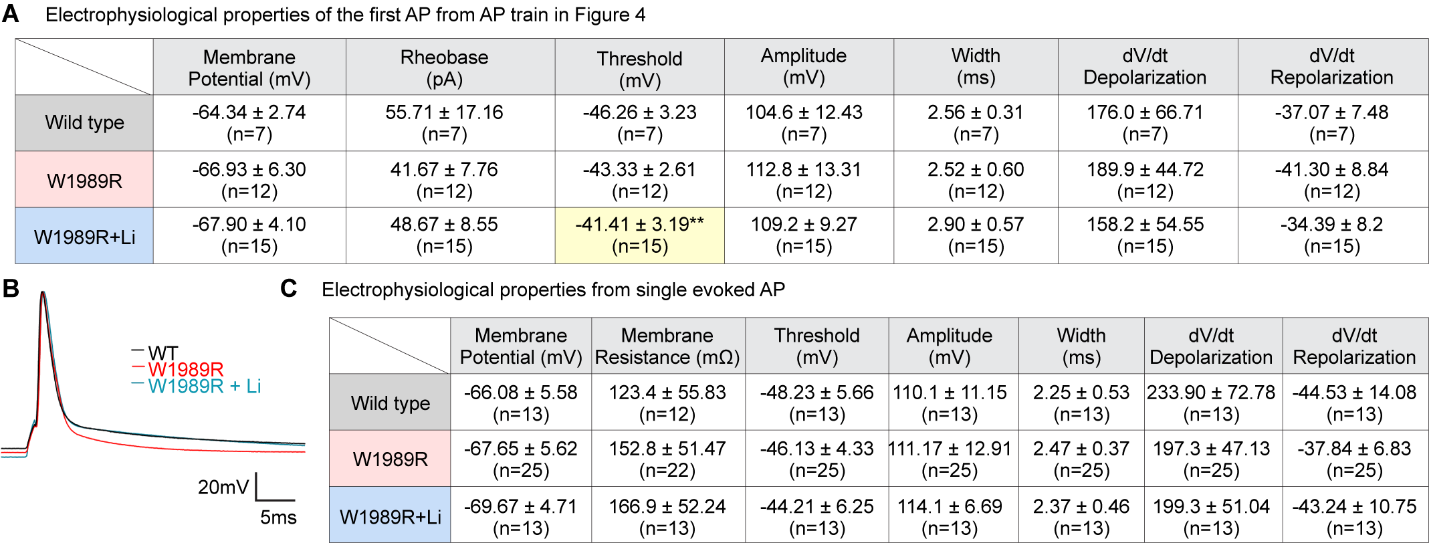

**Figure S4.**

**Effect of lithium treatment on single action potential (AP) properties.**

(A) Quantification of electrophysiological properties from the first AP in train shown in Figure 4 from layer II/III cortical pyramidal neurons in acute brain slices from wild-type (WT) and *Ank3* pW1989R homozygous mice with (W1989R+Li) or without lithium treatment (W1989R). Membrane potential: Ordinary one-way ANOVA (F (2, 31) = 1.308, P=0.2850). Rheobase: Ordinary one-way ANOVA (F (2, 31) = 0.3908, P=0.6798). Threshold: Ordinary one-way ANOVA (F (2, 31) = 6.251, P=0.0052). Tukey's multiple comparisons tests (**P= 0.0038 in WT vs. W1989R + Lithium). Amplitude: Ordinary one-way ANOVA (F (2, 31) = 1.152, P=0.3291). Width: Ordinary one-way ANOVA (F (2, 31) = 1.853, P=0.1738). dV/dt Depolarization: Ordinary one-way ANOVA (F (2, 31) = 1.157, P=0.3276). dV/dt Repolarization: Ordinary one-way ANOVA (F (2, 31) = 2.316, P=0.1155). (B) Representative single evoked APs from layer II/III cortical pyramidal neurons in acute brain slices from wild-type (WT) and *Ank3* pW1989R homozygous mice with (W1989R+Li) or without lithium treatment (W1989R). (C) Electrophysiological properties from evoked single APs. Membrane potential: Ordinary one-way ANOVA (F (2, 48) = 1.438, P=0.2474). Membrane resistance: Ordinary one-way ANOVA (F (2, 44) = 2.213, P=0.1214). Threshold: Ordinary one-way ANOVA (F (2, 48) = 1.933, P=0.1558). Amplitude: Ordinary one-way ANOVA (F (2, 48) = 0.4152, P=0.6626). Width: Ordinary one-way ANOVA (F (2, 48) = 1.098, P=0.3417). dV/dt Depolarization: Ordinary one-way ANOVA (F (2, 48) = 2.030, P=0.1424). dV/dt Repolarization: Ordinary one-way ANOVA (F (2, 48) = 2.340, P=0.1072). The ‘n’ represents the number of individual neurons per parameter in each condition. N=3-6 mice.
